# Unidirectional monosynaptic connections from auditory areas to the primary visual cortex in the marmoset monkey

**DOI:** 10.1101/373779

**Authors:** Piotr Majka, Marcello G. P. Rosa, Shi Bai, Jonathan M. Chan, Bing-Xing Huo, Natalia Jermakow, Meng K. Lin, Yeonsook S. Takahashi, Ianina H. Wolkowicz, Katrina H. Worthy, Ramesh Rajan, David H. Reser, Daniel K. Wójcik, Hideyuki Okano, Partha P. Mitra

**Affiliations:** Laboratory of Neuroinformatics, Nencki Institute of Experimental Biology, 02-093 Warsaw, Poland; Australian Research Council, Centre of Excellence for Integrative Brain Function, Monash University Node, Clayton, VIC 3800, Australia; Biomedicine Discovery Institute and Department of Physiology, Monash University, Clayton, VIC 3800, Australia; Laboratory for Marmoset Neural Architecture, RIKEN Center for Brain Science, Saitama 351-0106, Japan; Cold Spring Harbor Laboratory, Cold Spring Harbor, NY 11724, USA; School of Rural Health, Monash University, Churchill, VIC 3842, Australia; Department of Physiology, Keio University School of Medicine, Tokyo 160-8582, Japan

**Keywords:** striate cortex, auditory cortex, connections, primate, audiovisual integration

## Abstract

Until the late 20^th^ Century, it was believed that different sensory modalities were processed by largely independent pathways in the primate cortex, with cross-modal integration only occurring in specialized polysensory areas. This model was challenged by the finding that the peripheral representation of the primary visual cortex (V1) receives monosynaptic connections from areas of the auditory cortex in the macaque. However, auditory projections to V1 have not been reported in other primates. We investigated the existence of direct interconnections between V1 and auditory areas in the marmoset, a New World monkey. Labelled neurons in auditory cortex were observed following 4 out of 10 retrograde tracer injections involving V1. These projections to V1 originated in the caudal subdivisions of auditory cortex (primary auditory cortex, caudal belt and parabelt areas), and targeted parts of V1 that represent parafoveal and peripheral vision. Injections near the representation of the vertical meridian of the visual field labelled few or no cells in auditory cortex. We also placed 8 retrograde tracer injections involving core, belt and parabelt auditory areas, none of which revealed direct projections from V1. These results confirm the existence of a direct, nonreciprocal projection from auditory areas to V1 in a different primate species, which has evolved separately from the macaque for over 30 million years. The essential similarity of these observations between marmoset and macaque indicate that early-stage audiovisual integration is a shared characteristic of primate sensory processing.

## Introduction

The integration of visual and auditory information is an important process in daily life. Congruent stimulation can improve the detectability of sensory targets that are weak or masked by noise, especially in situations involving low signal to noise ratios (Witten and Knudsen 2005; Ross et al. 2006). The information from each modality appears to be weighted according to its estimated reliability: in audiovisual interactions vision tends to dominate spatial aspects of the combined percept, while audition dominates temporal aspects (Witten and Knudsen 2005; Burr and Alais 2006). Task difficulty also influences the likelihood of audiovisual fusion (Battaglia et al. 2003; Alsius et al., 2005; Wang et al. 2008).

The fact that information from one sensory modality can influence what we perceive through another modality implies the existence of polysensory neurons, which respond to more than one type of sensory input, or whose responses to one modality are enhanced or inhibited by concurrent stimulation of another modality. In the primate cerebral cortex, it has been known since the 1980s that overt polysensory responses occur in regions of the superior temporal, parietal and frontal cortices (see Beauchamp 2005; Macaluso and Driver 2005; van Atteveldt et al. 2014; Yau et al. 2015 for reviews). Yet, subtler cross-modal effects have been detected in regions originally thought to be strictly unimodal, including in visual and auditory areas (e.g. Ghazanfar and Schroeder 2006; Wang et al. 2008; Kayser et al. 2009).

In the classical view, the confluence of visual and auditory inputs only occurs in high-order, polysensory association cortices, after many hierarchical steps (Jones and Powell 1970; Felleman and Van Essen 1991). This anatomical model reflects the original physiological evidence that polysensory neurons are restricted to specialized association cortices (e.g. Benevento et al. 1977; Bruce et al. 1981). As these areas lack direct connections with primary and secondary sensory cortices, the model implies that low-level cross-modal physiological effects depend on chains of “feedback” connections. However, others have challenged this hierarchical model, highlighting the potential role of small projection pathways in providing “shortcuts” between early-stage unimodal processing areas (Schroeder et al. 2003; Cappe and Barone 2005, Palmer and Rosa 2006a; Smiley et al. 2007; Smiley and Falchier 2009; Falchier et al. 2010). One of the earliest, and most powerful findings underlying the current model of early-stage integration of visual and auditory information in the primate cortex refers to the existence of direct (monosynaptic) connections from auditory areas to the primary visual area (V1) in macaques. This projection (Falchier et al. 2002; Rockland and Ojima 2003) is reported to originate from caudal auditory areas, as well as adjacent polysensory cortex, and to target primarily the peripheral representation of V1.

Here we explored the existence of direct interconnections between auditory areas and V1 in the marmoset monkey (*Callithrix jacchus*), a New World monkey which is attracting increasing interest as a model species for studies of neural systems that are highly developed or specialized in the primate brain (Solomon and Rosa 2014; Mitchell and Leopold 2015; Majka et al. 2016; Miller et al. 2016; Eliades and Miller 2017; Hagan et al. 2017; Oikonomidis et al. 2017; Okano et al. 2016; Okano and Kishi 2017; Silva 2017). Previous work in this species has indicated that sparse projections from auditory cortex target the middle temporal visual area (MT; Palmer and Rosa 2006a), but studies of V1 connections have not reported afferents from the auditory cortex (Rosa and Tweedale 2000; Lyon and Kaas 2001). Here we report that the peripheral representation of V1 receives a sparse projection from the caudal auditory cortex and adjacent polysensory areas, which is not reciprocated by a V1 projection to auditory cortex. The similarity of observations between marmoset and macaque, species that have evolved separately for over 30 million years, indicate that early-stage audiovisual integration is a shared characteristic of primate sensory processing.

## Materials and Methods

Experiments involved 11 adult marmoset monkeys (*Callithrix jacchus*). Tracer injections were performed both at Monash University (cases CJ64, CJ75, CJ82, CJ122, CJ174, CJ178, CJ180 and CJ802) and the RIKEN Brain Science Institute (BSI, which was re-established as RIKEN Center for Brain Science [CBS] in April 2018); cases M820, M822 and M1146). Very similar methods where used on both locations (where applicable, differences are detailed below). Experiments in Australia conformed to the Australian Code of Practice for the Care and Use of Animals for Scientific Purposes, and were approved by the Monash University Animal Experimentation Ethics Committee, and those in Japan were approved by the Institutional Animal Care and Use Committee at RIKEN, and conducted in accordance with the Guidelines for Conducting Animal Experiments at RIKEN BSI. Participation of Australian Researchers in surgeries conducted at the RIKEN BSI was authorized by a field license obtained from the Monash University Animal Experimentation Ethics Committee. Each of the animals received multiple injections of fluorescent tracers, which were aimed at specific locations in the cortex using stereotaxic coordinates (Paxinos et al., 2012). The exact areas involved in each injection were subsequently determined based on histological examination (Table 1). A full list of the abbreviations of cortical areas that contained labelled neurons following the present injections is presented in Table 2. In all figures the sections and maps are illustrated using the convention appropriate for a left hemisphere, to facilitate comparisons between cases and with earlier publications from our laboratories. The actual hemisphere in which the injections were placed is given for each case in Table 1. The extents of diamidino yellow (DY) and fast blue (FB) injection sites were estimated according to the criteria defined by Condé (1987). Other injections were estimated as the volume of cortex containing fluorescent dye in the extracellular space (which is limited to the neighbourhood of the needle track; Schmued et al., 1990).

**Table 1:** Animals and injections.

| Animal | Age (months) | Sex | Weight (g) | Hemisphere injected | Survival time (days) | Tracer | Area(s) injected | Layers injected | Injection volume (mm<sup>3</sup>)<sup>a</sup> |
| --- | --- | --- | --- | --- | --- | --- | --- | --- | --- |
| CJ82 | 28 | F | 375 | L | 15 | FE | V1 | 3 | 0.14 |
|  |  |  |  |  |  | FR | V1 | 2-6 | 0.10 |
|  |  |  |  |  |  | DY | V2, V1 | 1-3 | 0.11 |
| CJ174 | 30 | F | 342 | R | 17 | CTB (g) | V1 | 1-3 | 0.11 |
|  |  |  |  |  |  | CTB (r) | V1 | 1-3 | 0.03 |
|  |  |  |  |  |  | FB | V1 | 1-6 | 1.19 |
| CJ178 | 26 | F | 344 | R | 18 | FB | V1 | 1-6 | 0.49 |
| M820 | 83 | F | 400 | L | 30 | FB | V1 | 1-6 | 0.63 |
| M822 | 85 | F | 355 | L | 29 | FB | V1, V2 | 1-6 | 0.81 |
| M1146 | 85 | F | 394 | L | 21 | FB | V1 | 1-3 | 0.05 |
| CJ64 | 16 | M | 303 | R | 14 | FR | CM, A1 | 1-5 | 0.17 |
|  |  |  |  |  |  | FE | AL, ML | 1-5 | 0.17 |
| CJ75 | 16 | M | 356 | R | 10 | FR | A1 | 1-6 | 0.12 |
| CJ122 | 34 | M | 318 | R | 14 | DY | RT, AL | 1-6 | 0.17 |
|  |  |  |  |  |  | FR | CPB, AL | 1-4 | 0.05 |
| CJ180 | 32 | M | 374 | R | 11 | FB | RT, AL, RTL | 1-6 | 0.99 |
|  |  |  |  |  |  | CTB (g) | ML, CPB | 1-6 | 0.11 |
| CJ802 | 43 | M | 352 | R | 22 | CTB (g) | ML, CL, CPB | 1-6 | 0.17 |

<sup>a</sup> Injection site volume estimated across multiple histological sections, using the criteria proposed by Condé (1987) and Schmued et al. (1990). Measurements uncorrected for shrinkage due to histological processing.

**Table 2:** Abbreviations of names of cortical areas.

|  |  |
| --- | --- |
| 1/2 | Areas 1 and 2 of somatosensory cortex |
| 6DR | Cytoarchitectural area 6 of cortex, dorsorostral part |
| 8aV | Cytoarchitectural area 8a of cortex, ventral part |
| 8b | Cytoarchitectural area 8b of cortex |
| 10 | Cytoarchitectural area 10 of cortex (frontopolar cortex) |
| 11 | Cytoarchitectural area 11 of cortex |
| 12L | Cytoarchitectural area 12 of cortex (area 47), lateral part |
| 12M | Cytoarchitectural area 12 of cortex (area 47), medial part |
| 13L | Cytoarchitectural area 13 of cortex, lateral part |
| 13M | Cytoarchitectural area 13 of cortex, medial part |
| 19M | Cytoarchitectural area 19 of visual cortex, medial part |
| 23a | Cytoarchitectural area 23a of cortex |
| 23V | Cytoarchitectural area 23 of cortex, ventral part |
| 32 | Cytoarchitectural area 32 of cortex |
| 36 | Cytoarchitectural area 36 of cortex |
| 45 | Cytoarchitectural area 45 of cortex |
| A1 | Primary auditory area |
| AL | Anterolateral auditory area |
| CL | Caudolateral auditory area |
| CM | Caudomedial auditory area |
| CPB | Caudal parabelt auditory area |
| DM | dorsomedial visual area (V6) |
| FST | Fundus of superior temporal sulcus area |
| GI | Granular insular cortex |
| LIP | Lateral intraparietal area |
| ML | Middle lateral auditory area |
| MST | Medial superior temporal area |
| MT | Middle temporal area (V5) |
| MTc | Middle temporal crescent area (V4T) |
| PaIM | Parainsular cortex, medial part |
| PFG | Cytoarchitectural area PFG |
| PG | Cytoarchitectural area PG |
| PGa/IPa | Cytoarchitectural areas PGa and IPa |
| PR | Parietal rostral area (somatosensory) |
| ProSt | Area prostriata |
| PV | Parietal ventral area (somatosensory) |
| R | Rostral auditory area |
| ReI | Retroinsular area |
| RM | Rostromedial auditory area |
| RPB | Rostral parabelt auditory area |
| RT | Rostrotemporal auditory area |
| RTL | Rostrotemporal lateral auditory area |
| S2 | Secondary somatosensory cortex |
| TE1 | Cytoarchitectural area TE, part 1 |
| TE3 | Cytoarchitectural area TE, part 3 |
| TEO | Cytoarchitectural area TE, occipital transition part |
| TF | Cytoarchitectural area TF |
| TFO | Cytoarchitectural area TF, occipital transition part |
| TH | Cytoarchitectural area TH |
| TL | Cytoarchitectural area TL |
| TLO | Cytoarchitectural area TL, occipital transition part |
| TPO | Temporo-parieto-occipital association area (superior temporal polysensory cortex, STP) |
| TPOc | Temporo-parieto-occipital association area, caudal subdivision |
| TPPro | Temporopolar proisocortex |
| TPt | Temporoparietal transitional area |
| V1 | Primary visual cortex |
| V2 | Visual area 2 |
| V3 | Visual area 3 (ventrolateral posterior area, VLP) |
| V3a | Visual area 3a (dorsoanterior area, DA) |
| V4 | Visual area 4 (ventrolateral anterior area, VLA) |
| V6a | Visual area 6a (posterior parietal medial area) |
| VIP | Ventral intraparietal area |

### Surgeries

The surgical procedures have been described in detail previously (Reser et al. 2013; Burman et al., 2014a, b). Intramuscular (i.m.) injections of atropine (0.2 mg/kg) and diazepam (2 mg/kg) were administered as premedication, before each animal was anaesthetized with alfaxalone (10 mg/kg, i.m.) 30 min later. Dexamethasone (0.3 mg/kg, i.m.) and amoxicillin (50 mg/kg, i.m.) were also administered prior to positioning the animals in a stereotaxic frame. Body temperature, heart rate, and blood oxygenation (PO2) were continually monitored during surgery, and when necessary, supplemental doses of anaesthetic were administered to maintain areflexia. Incisions of the dura mater were made over the intended injection sites to limit exposure of the brain’s surface. Tracer injections were placed in the same hemisphere in each animal.

Six types of fluorescent tracers were used (Table 1): fluororuby (FR; dextran-conjugated tetramethylrhodamine, molecular weight 10,000, 15%), fluoroemerald (FE; dextran-conjugated fluorescein, molecular weight 10,000, 15%), fast blue (FB), diamidino yellow (DY) and CTB (cholera toxin subunit B, conjugated with either Alexa 488 [CTBg] or Alexa 594 [CTBr]). The dextran tracers resulted in bidirectional transport, but only retrograde labelling is reported here. In most cases the tracers were injected using 25 μl constant rate microsyringes (Hamilton, Reno, NV) fitted with a fine glass micropipette tip. In cases M820, M822 and M1146, the injections used a Nanoject II injector (Drummond Scientific, Broomall, PA), also fitted with a micropipette, with dosage controlled by Micro4 microsyringe pump controller (World Precision Instruments, Sarasota, FL). Each tracer was injected over 15–20 min, with small deposits of tracer made at different depths. Following the last deposit, the pipette was left in place for 3–5 min to minimize tracer reflux. Estimates of injection extent for each case, drawn under microscopic examination, are listed in Table 1. After the injections, the surface of the brain was covered with moistened ophthalmic film, over which the dural flaps were carefully arranged. The excised bone fragment was repositioned and secured in place with dental acrylic, and the wound closed in anatomical layers. Postoperative injectable analgesics were administered immediately after the animal exhibited spontaneous movements (Temgesic 0.01 mg/kg, i.m., and Carprofen 4 mg/kg, s.c.), followed by oral Metacam (0.05 mg/kg) for 3 consecutive days.

### Tissue processing and data analysis (experiments performed at Monash University)

Survival times were between 11 and 22 days (Table 1), after which the animals were anesthetized with alfaxalone (10 mg/ml i.m.) and, following loss of consciousness, administered an overdose of sodium pentabarbitone (100 mg/kg, i.v.). They were then immediately perfused through the heart with 1 l of heparinized saline, followed by 1 l of 4% paraformaldehyde (PFA) in 0.1 M phosphate buffered saline (PBS; pH 7.4). The brains were post-fixed in the same medium for at least 24 hours, and then immersed in buffered paraformaldehyde with increasing concentrations of sucrose (10–30%). They were then sectioned (40 μm thickness) in the coronal plane, using a cryostat. One section in five was mounted unstained for examination of fluorescent tracers, and coverslipped after quick dehydration (2× 100% ethanol) and defatting (2× xylene). Adjacent sections were stained for Nissl substance, cytochrome oxidase, and myelin, following standard protocols (Gallyas, 1979; Wong-Riley, 1979). The remaining section in each series was stored in cryoprotectant solution in a freezer, to be used as a backup in case of unsatisfactory staining or damage during processing of the histological sections. Hence, the spacing between adjacent sections in each series was 200 μm in these cases.

Sections were examined using a Zeiss Axioplan 2 epifluorescence microscope. Labelled neurons were identified using ×10 or ×20 dry objectives, and their locations within the cortex and subcortical structures were mapped using a digitizing system (MD Plot3, Accustage) attached to the microscope. To minimize the problem of overestimating the number of neurons due to inclusion of cytoplasmic fragments, labelled cells were accepted as valid only if a nucleus could be discerned. This was straightforward in the case of DY, since this tracer only labels the neuron’s nucleus (Keizer et al., 1983). In the case of tracers that label the cytoplasm (FB, FE, FR, CTBg and CTBr), the nucleus was discerned as a profile in the centre of a brightly lit, well-defined cell body, which in the vast majority of cases had an unmistakable pyramidal morphology.

To allow assessment of the areas in which injection sites and labelled neurons were located, each brain was reconstructed in 3 dimensions. Nissl-stained sections were scanned using Aperio Scanscope AT Turbo (Leica Biosystems) at 20× magnification, providing a resolution of 0.50 μm/pixel. Within each coronal section, we identified the cortical areas proposed by Paxinos et al. (2012), based primarily on cytoarchitectural features observable in Nissl-stained sections, but also using differences in myeloarchitecture and intensity of cytochrome staining as additional criteria to refine the delineations (e.g.Rosa et al. 2005, 2009; Palmer and Rosa 2006a, b; Burman and Rosa 2009; Burman et al. 2006, 2014a, b, 2015). This was followed by computerized 3-dimensional reconstruction of the entire hemisphere, and registration of each case to a template brain (Majka et al. 2016), to allow comparison between results from different cases. The registration procedure was supported by manually-drawn cyto-, myelo- and chemoarchitectonic boundaries, thus ensuring that the spatial relationships between the injection sites, labelled neurons, and histological boundaries were adequately maintained.

### Tissue processing and data analysis (experiments performed at RIKEN)

The concurrent use of virus-based anterograde tracers (results to be reported separately) demanded longer survival times (21–30 days; Table 1) for these animals. At the end of the survival time they were anesthetized with a combination of ketamine (10 mg/kg) and diazepam (2 mg/kg). Following the loss of consciousness, they were administered a lethal injection of pentobarbital (80 mg/kg) and perfused through the heart with 500 ml of heparinised PBS (pH 7.2) and 500 ml of 4% PFA in 0.1 M phosphate buffer (PB). The brains were removed from the skull, stored in the same fixation medium overnight, and then transferred to a 0.1 M PB solution. A post-mortem MRI (data not used in the present study) was obtained at this stage, with Diffusion Tensor Imaging (DTI) and 300 μm T2-weighted images (T2WI). Once the ex-vivo MRI was finished, the brains were transferred into increasing concentrations of sucrose in 0.1 M PB (10% for 24 hours, then 30% until sunk), and then embedded in Neg50 frozen section medium (Richard Allen Scientific, Waltham, MA) using a custom made polylactic acid base, a 3d-printed brain mould, and 2-methylbutane chilled with dry ice. Following embedding the brains were stored at −80°C until sectioning.

Each brain was sectioned at 20 μm thickness in the coronal plane using a cryostat and a tape transfer technique (Pinsky et al. 2015), which minimises distortions in section morphology during mounting. Three consecutive series of sections were stained (for Nissl substance, myelinated fibres (Gallyas, 1979), and cholera toxin subunit B immunocytochemistry (Angelucci et al., 1996; results not reported here), whereas the fourth series was kept unstained for fluorescent tracer analysis. Thus, the spacing between consecutive sections in each series was 80 μm in these cases. Following curing with UV light, the slides used for the analysis of fluorescence were placed in xylene and coverslipped. All slides were scanned using a Nanozoomer 2.0 HT (Hamamatsu, Japan) equipped with a 20× objective (0.46 μm/pixel in plane) at 12-bit depth. The Nissl and myelin series were scanned with bright field illumination, and the fluorescence series with a tri-pass filter cube (FITC/TX-RED/DAPI) for excitation. All imaging data were processed with a high-throughput neurohistological pipeline (Lin et al. 2018), resulting in high resolution images (e.g. Fig. 5).

## Results

Figure 1 illustrates the locations of the centres of the 18 tracer injections analysed in the present study. Of these, 10 were located in V1, and 8 in various auditory areas (Table 1). Although the present analysis is focused on the evidence for direct cross-modal (auditory-visual) cortico-cortical connections, each of the injections also resulted, as expected, in retrograde label that extended across multiple cortical and subcortical areas. The full pattern of corticocortical label obtained in 8 of the animals (experiments conducted at Monash University) can be visualised, section by section, in a freely accessible web site (http://www.marmosetbrain.org/), which includes online tools for quantification; here, it will be summarised using “unfolded” representations of the cortex obtained by computational registration of the areas in each brain to a common template (Majka et al. 2016), and only direct connections between visual and auditory areas will be discussed. Data from the other 3 animals (experiments conducted at the RIKEN CBS) are hosted in http://riken.marmoset.brainarchitecture.org.

**Figure 1:**
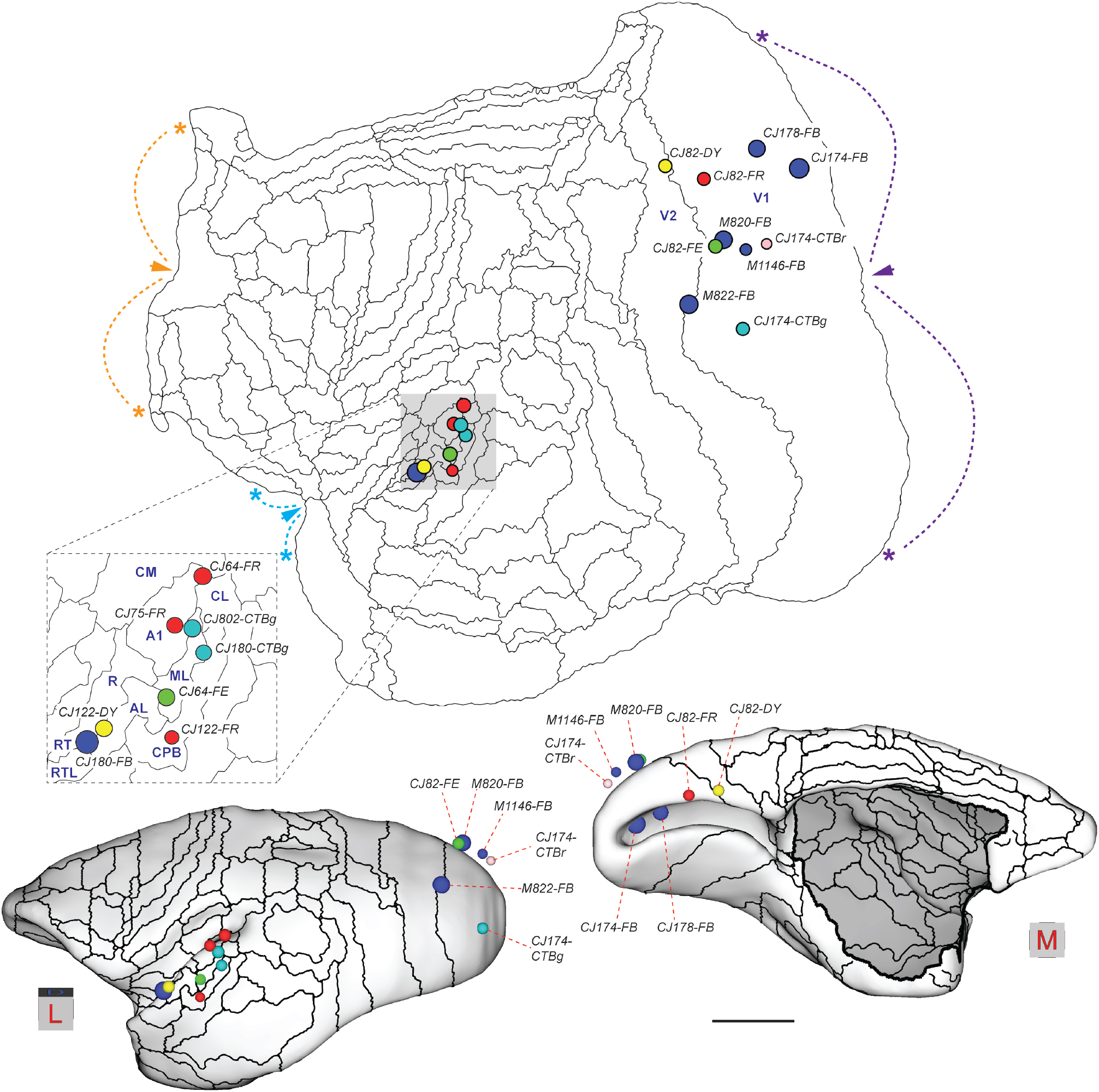
Locations of 18 tracer injections analysed in the present study, plotted on a template of the marmoset cortex (reconstructed by Majka et al. 2016, from data published by Paxinos et al. 2012). The template can be freely downloaded from http://www.marmosetbrain.org/reference. **Top**: Unfolded reconstruction of the cortex (left hemisphere representation; rostral to the left, medial to the top) showing the location of each injection registered to the template. The colours of each circle represent the tracer used (blue- FB; yellow- DY; red- FR; green- FE; pink- CTBr; teal- CTBg), and their radius is proportional to that of a sphere with equal volume to the injection site reconstructed across histological sections (Table 1). The boundaries of cortical areas are indicated by thin lines. The auditory cortex (shaded in grey) is shown magnified in the insert. Arrowheads and asterisks indicate three points along the perimeter of the map where discontinuities were made to minimise distortions (fundus of the calcarine sulcus in purple, boundary between orbitofrontal cortex and medial frontal cortex in orange, piriform cortex in blue). For abbreviations, see Table 2. **Bottom**: 3-dimensional views of the marmoset brain from dorsolateral (left) and medial (right) perspectives, showing the locations of the injections in a reconstruction of the cortex at the level of its mid- thickness (for this reason, injections located in the supragranular layers appear to “float” above the cortex). Scale bar: 5 mm.

### Projections from auditory areas to V1

Our main observation is that the caudal auditory cortex sends a sparse projection to the portions of V1 located in the calcarine sulcus and midline. The most obvious label in auditory areas was observed in animal CJ174, following an injection of FB in the dorsal bank of the calcarine sulcus, with slight invasion of the ventral bank. This injection was centred in the lower quadrant representation near the horizontal meridian, at approximately 15-20^°^ eccentricity (Fritsches and Rosa 1996; Chaplin et al. 2013), but also included a portion of the upper quadrant representation (Fig. 2A). This resulted in widespread intrinsic label in V1, and extrinsic label in the peripheral representations of both dorsal and ventral visual cortex areas (Fig. 2A). There was no evidence of direct tracer spread to other areas (Fig. 2B, C). Figure 2D shows a summary view of the cortico-cortical connections revealed by this injection, which yielded labelled neurons in the various auditory and polysensory areas, and Figure 3 illustrates examples of the locations of labelled neurons. V1-projecting neurons were found in the primary auditory area (A1; 22 neurons), the caudomedial (CM), caudolateral (CL) and middle lateral (ML) belt areas (30, 21 and 19 neurons, respectively), the caudal parabelt area (CPB; 7 neurons) and in the temporoparietal transition auditory association area (TPt; 15 neurons). There were also labelled neurons in putative polysensory areas (TPO and PGa/IPa). Despite the clearly defined labelled cells, the combined auditory projection accounted for a small proportion of the afferents to V1 (114 / 15,148, or 0.75% of the labelled cortico-cortical extrinsic connection neurons). Most (>90%) of the labelled neurons were located in the infragranular layers, although in some sections supragranular neurons were also observed (Fig. 3).

**Figure 2:**
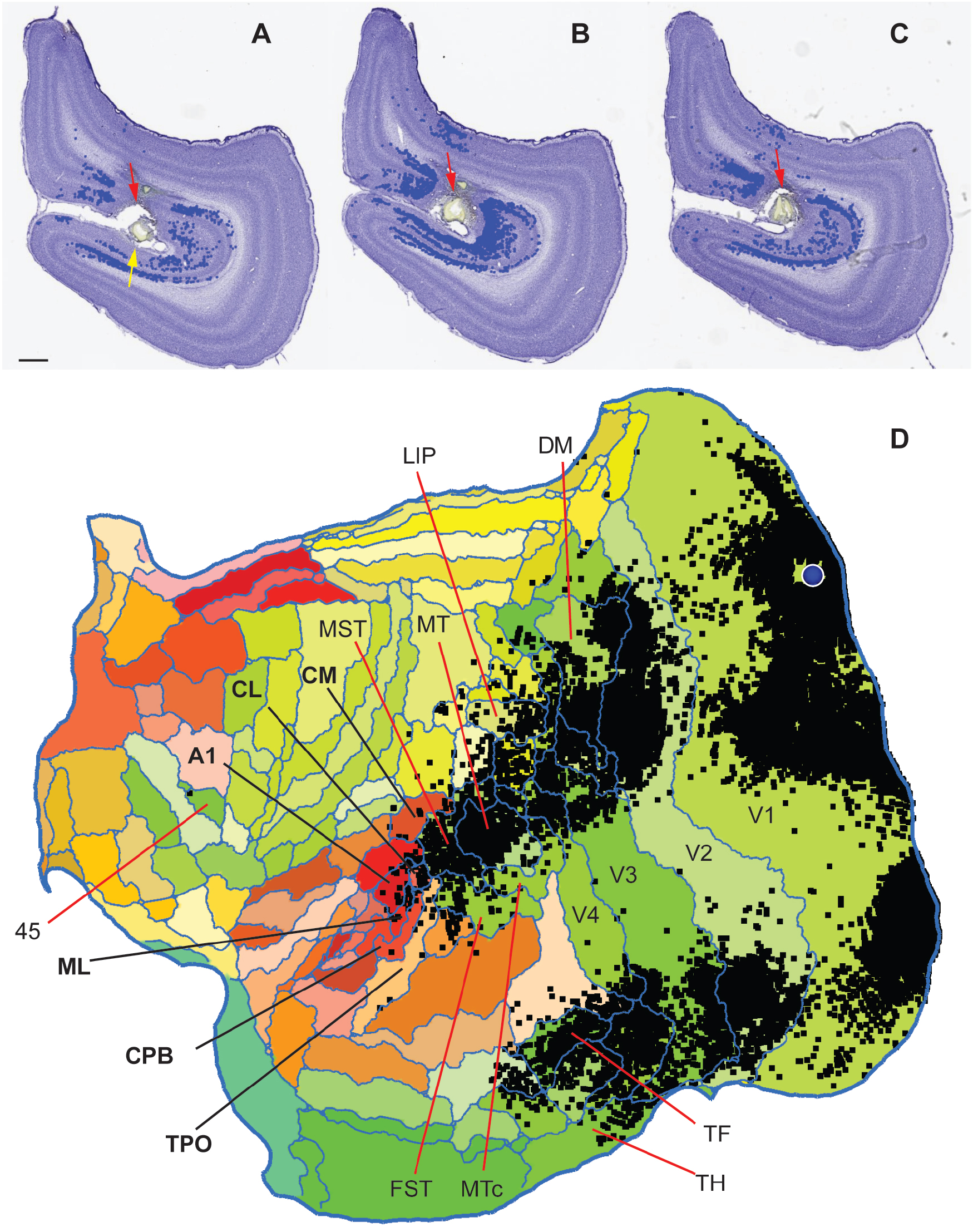
Summary of results of a FB injection in animal CJ174. **A-C**: Extent of the injection site, in coronal sections through the right hemisphere of a marmoset brain (anteroposterior [AP] levels −8.6 mm to −9.0 mm; section A is the most rostral; lateral to the right). Scale bar = 1 mm. The red arrows point to the main injection site in the dorsal bank of the calcarine sulcus, and the yellow arrow points to the slight invasion of the ventral bank. At these levels the entire section is formed by V1. Blue symbols represent labelled neurons forming intrinsic connections within V1. **D**: Unfolded reconstruction of the cortex showing the locations of neurons labelled by the tracer injection (black squares) following registration to the template. The injection site is indicated by the blue circle. In this and following figures, “flat” maps are oriented in the convention adopted for the Marmoset Brain Architecture Project (http://marmosetbrain.org), which represent a left hemisphere (rostral to the left, medial to the top), to facilitate comparison across cases (e.g. Paxinos et al. 2012; Majka et al. 2016), and the auditory cortex corresponds to a cluster of areas indicated in tones of red and orange. For the actual hemisphere injected, see Table 1. For orientation, the locations of several cortical areas are indicated.

**Figure 3:**
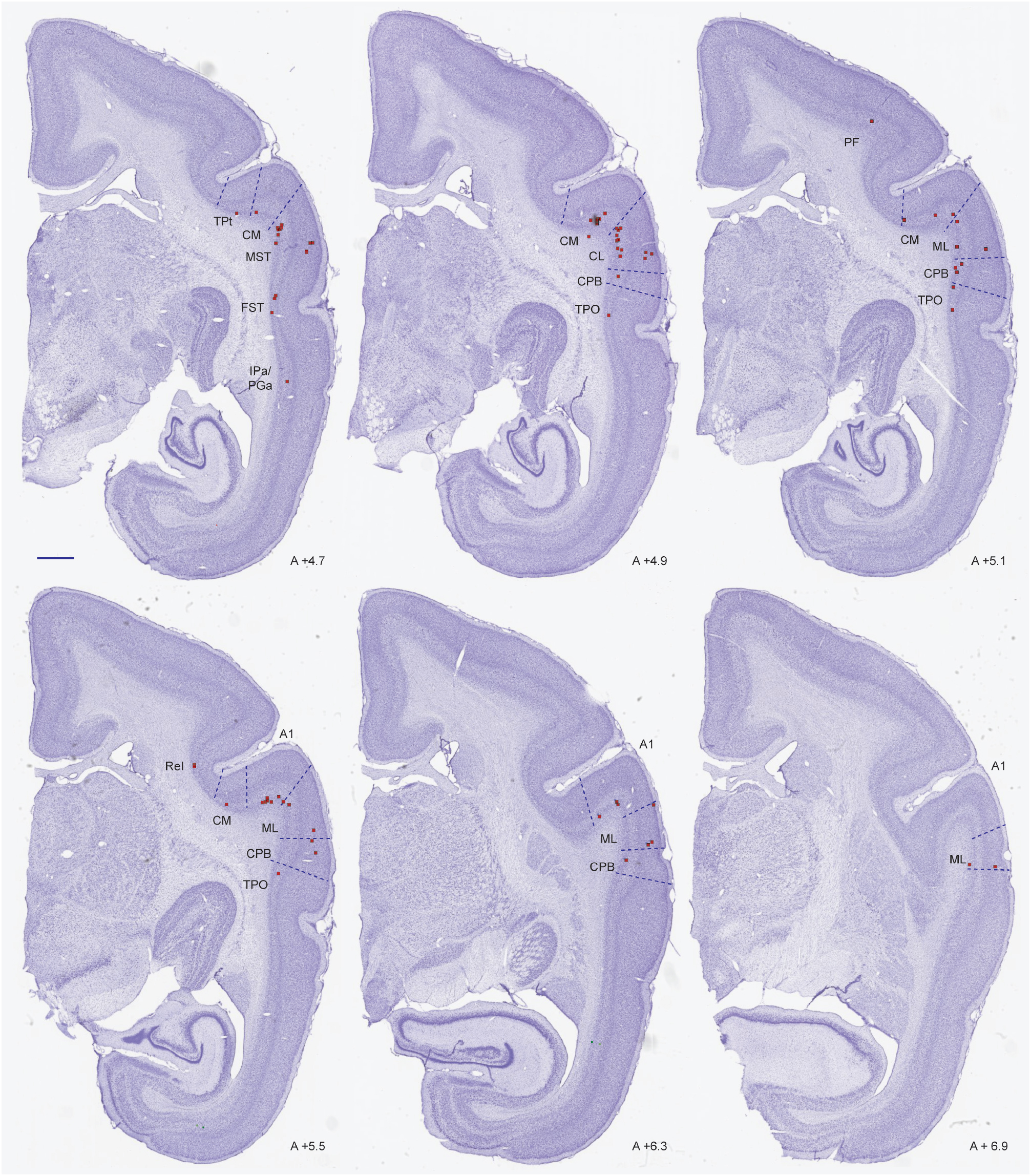
Coronal sections through auditory cortex in case CJ174-FB, showing the locations of some of the neurons labelled by the V1 injection shown in Figure 2 (red squares). The AP level of each section is indicated. Scale bar (top left): 1 mm.

Figure 4 illustrates data from a second animal (CJ178) with a FB injection in the calcarine sulcus. In this case, damage subsequent to the penetration of the injection syringe resulted in necrosis of the overlying optic radiations and parts of V2, which made impossible to ascertain the possibility of leakage (Fig. 4A). Nonetheless, the pattern of label in the cortex (Fig. 4B) was highly compatible with that observed in CJ174, although the total number of labelled neurons was much smaller (extrinsic cortico-cortical neurons in CJ178: 3,837). Based on comparison with Chaplin et al. (2013), this injection site was located further away from the representation of the horizontal meridian, in a region of V1 corresponding to 25–30^°^ eccentricity in the lower visual field. Label in auditory cortex was again sparse (0.23% of the extrinsic projections to V1), but originated in a subset of the same areas that formed projections in CJ174.

**Figure 4:**
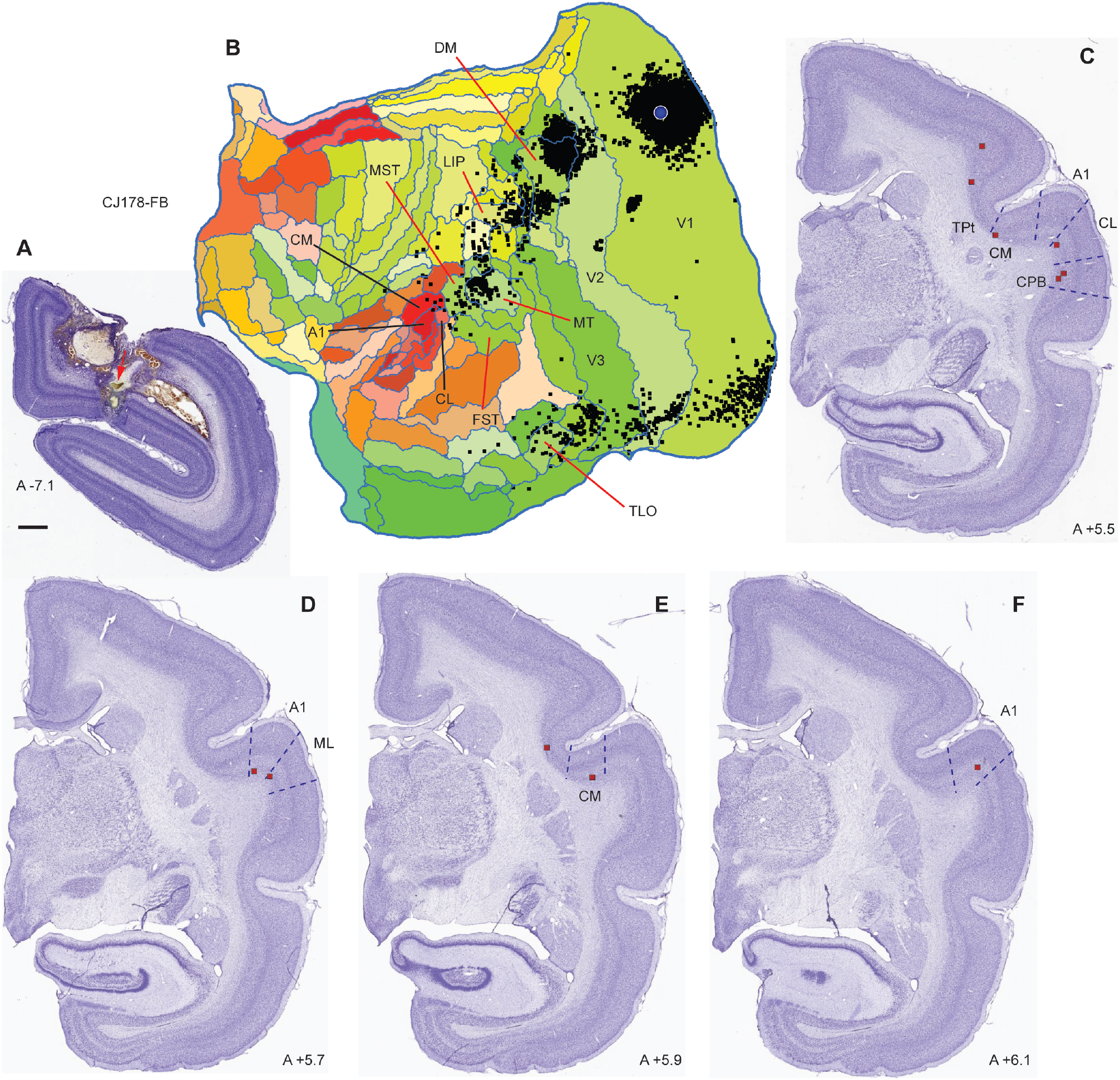
Summary of results in case CJ178-FB. **A:** coronal section through the centre of the FB injection site. In this case there was damage to the cortex overlying the calcarine sulcus subsequent to the injection. The injection site is indicated by the red arrow. Scale bar= 1 mm. B: Unfolded reconstruction of the cortex showing the locations of neurons labelled by the tracer injection (Black squares) following registration to the template. The injection site is indicated by the blue circle. C: Examples of coronal sections containing labelled neurons in auditory cortex, with their locations indicated by red squares. Conventions as in Figures 2 and 3.

Figure 5 illustrates examples of FB-labelled neurons in auditory areas following the injection in case M820. The injection site was located in the representation of parafoveal vision in V1, close to the midline, and away from the representation of the vertical meridian (Figs. 1, 5A), as confirmed by the distribution of label in the lateral geniculate nucleus (Fig. 5B). Labelled neurons were located in the same areas that formed projections to V1 in cases CJ174 and CJ178 (e.g. Fig. 5C, Fig. 6). Quantification of these data confirmed the existence of a strong laminar bias in the projection, whereby each of the labelled cells was located in the infragranular layers. A fourth case, M822- FB, had an injection site at the V1/ V2 border, at a more central location (~3° eccentricity). Only a few labelled neurons were observed in auditory areas (n=6), all of which were located in the infragranular layers of caudal lateral belt cortex.

**Figure 5:**
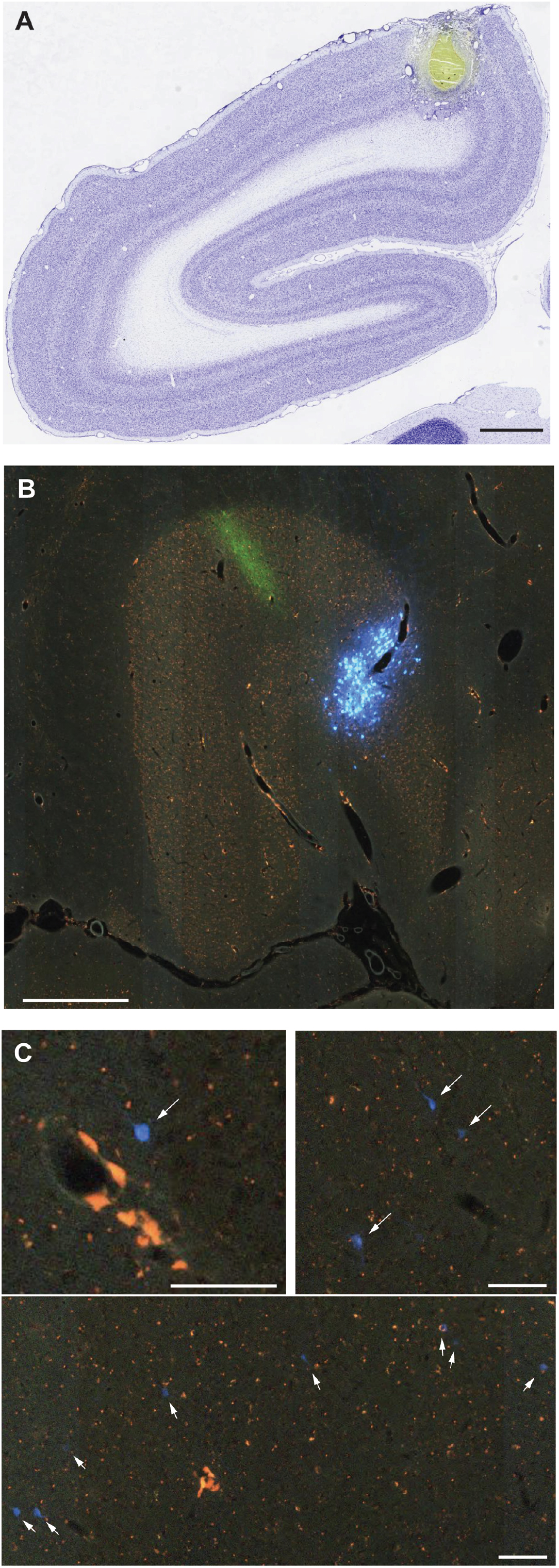
Results from case M820. **A:** Location of the FB injection site in V1, visualised in a Nissl stained section. Scale bar = 1 mm. **B:** Pattern of label in the lateral geniculate nucleus, confirming placement in V1, away from the representation of the vertical meridian (White et al. 1998). Label resulting from an injection of anterograde tracer in V1 is also visible (green). Scale bar = 500 μm. **C:** Example FB-labelled neurons (arrows) in auditory areas, visualised in digital images obtained at various magnifications (scale bars = 20 μm). The top left image shows a single cell in caudal parabelt cortex, the top right panel shows a small cluster of labelled neurons in area CL, and the bottom panel illustrates a line of layer 6 cells crossing the border between CL and CM.

**Figure 6:**
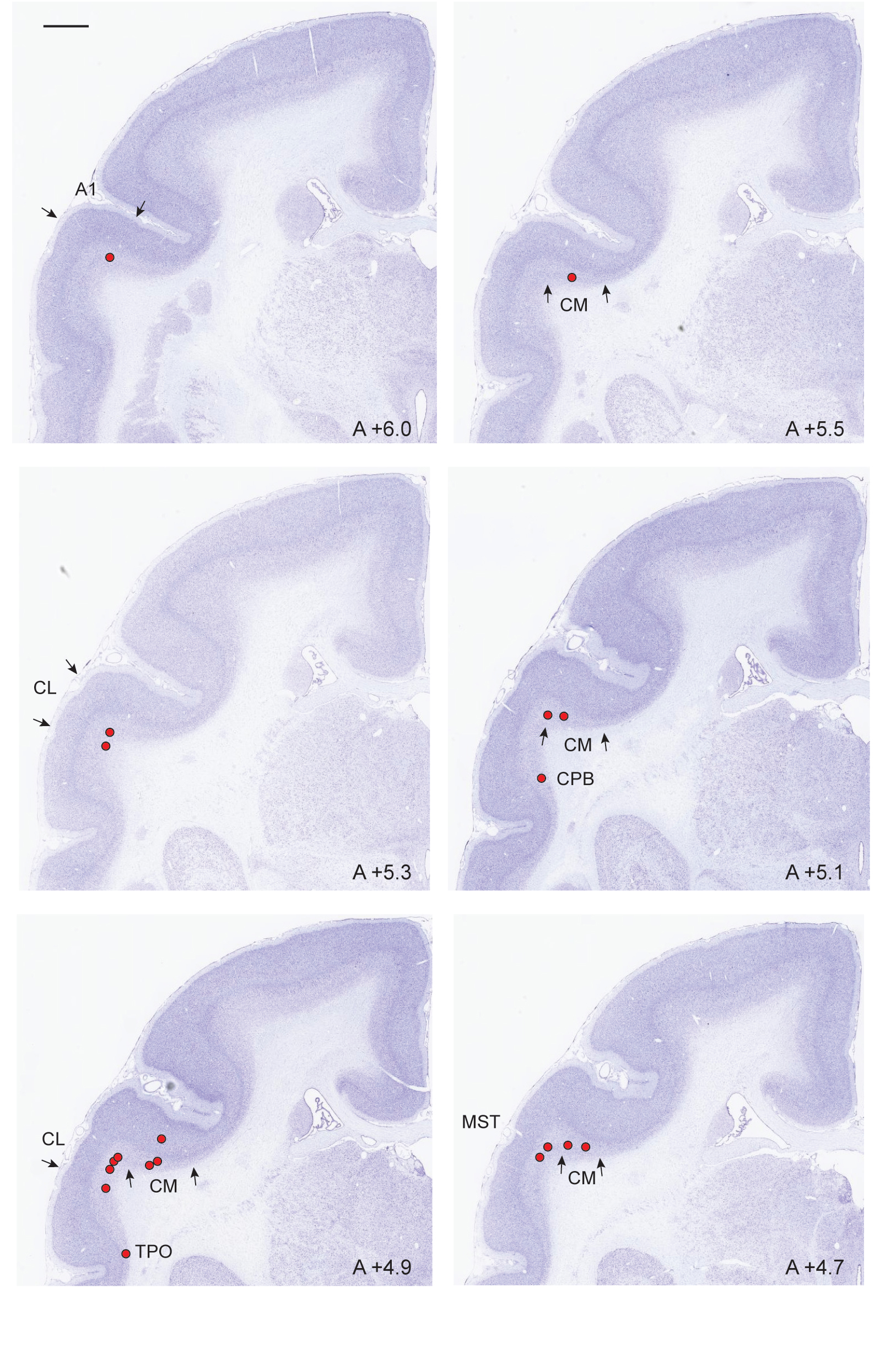
Examples of the locations of labelled neurons in auditory and adjacent areas in case M820. Scale bar (top left): 1 mm.

### Injections that revealed no evidence of projections from the auditory cortex

Six other injections showed no evidence of labelled neurons in auditory areas, despite clear evidence of long-range tracer transport over long distances (e.g, to the frontal lobe). Summary maps of the pattern of label in five of these cases are shown in Figure 7 (the results from animal M1146 could not be registered to the template, preventing display in flat map format). Four of the injections that resulted in no label in auditory cortex (CJ174-CTBr, CJ174-CTBg, CJ82-FE and M1146) were in parts of V1 representing central vision (<5^°^ eccentricity), and two (CJ82-FR and CJ82-DY) were located at the representation of the peripheral lower vertical meridian of the visual field (~10° and 20° eccentricity, respectively) in V1 (CJ82-FR) or V2, with small involvement of V1 (CJ82-DY). With one exception (CJ82-FR) these injections were all restricted to the supragranular layers; hence they may not have labelled the entire complement of afferents to this area.

**Figure 7:**
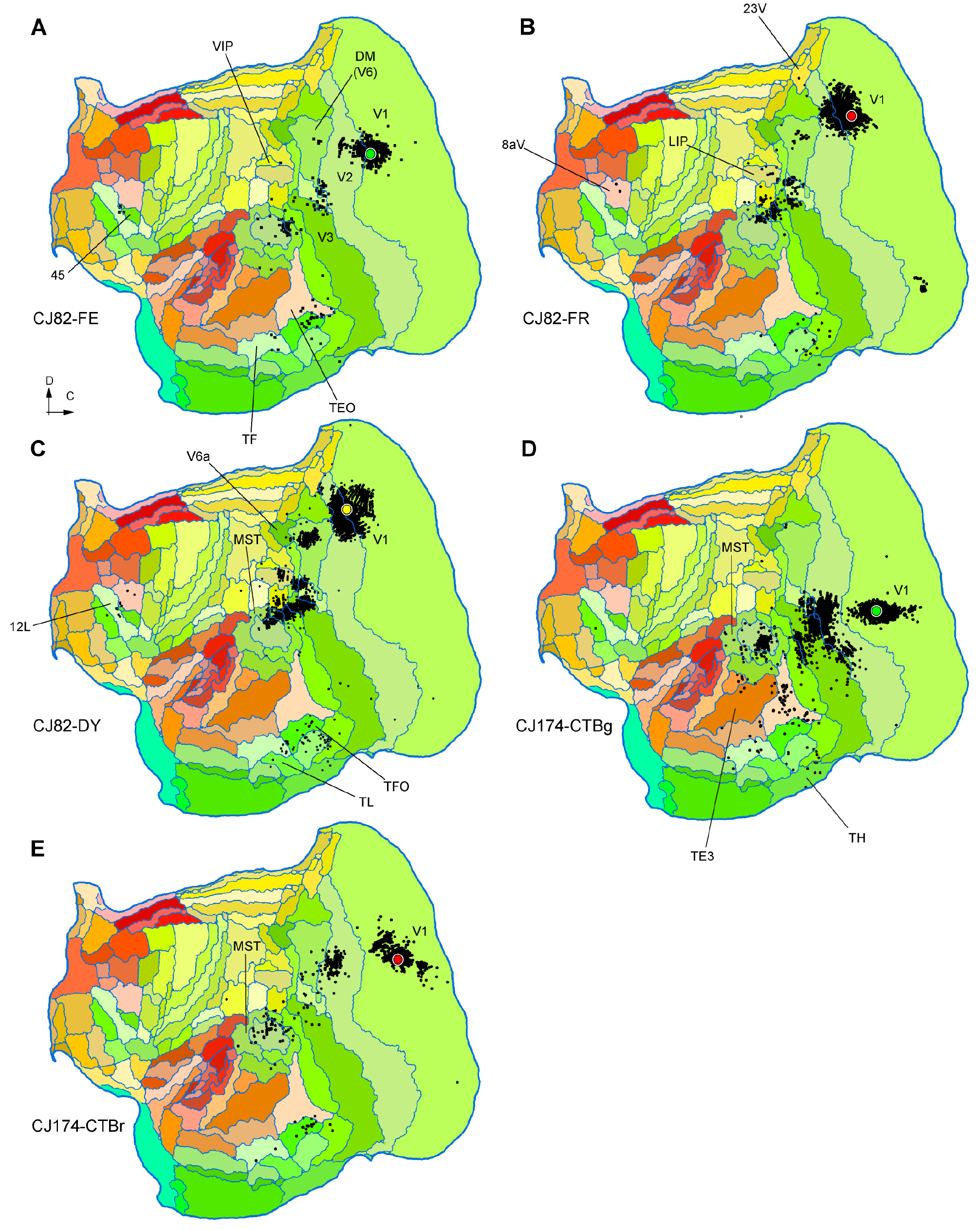
**A-B:** Unfolded reconstructions of the cortex showing the locations of labelled neurons following tracer injections which did not result in any labelled neurons in auditory cortex. The cases illustrated in A, B, D and F correspond to injections that were entirely contained within V1, while the one illustrated in C was centred in V2, with slight V1 invasion. Conventions as in Fig. 2. Several cortical areas where long-range projection neurons were labelled are indicated for orientation. For abbreviations, see Table 2.

### Injections in auditory cortex

To explore the possibility of a reciprocal projection from auditory areas to V1, we have studied the connections revealed by injections in various auditory areas. As detailed below, despite label in a few other visual areas, none of these revealed direct projections from V1 (Figs. 8 and 9).

**Figure 8:**
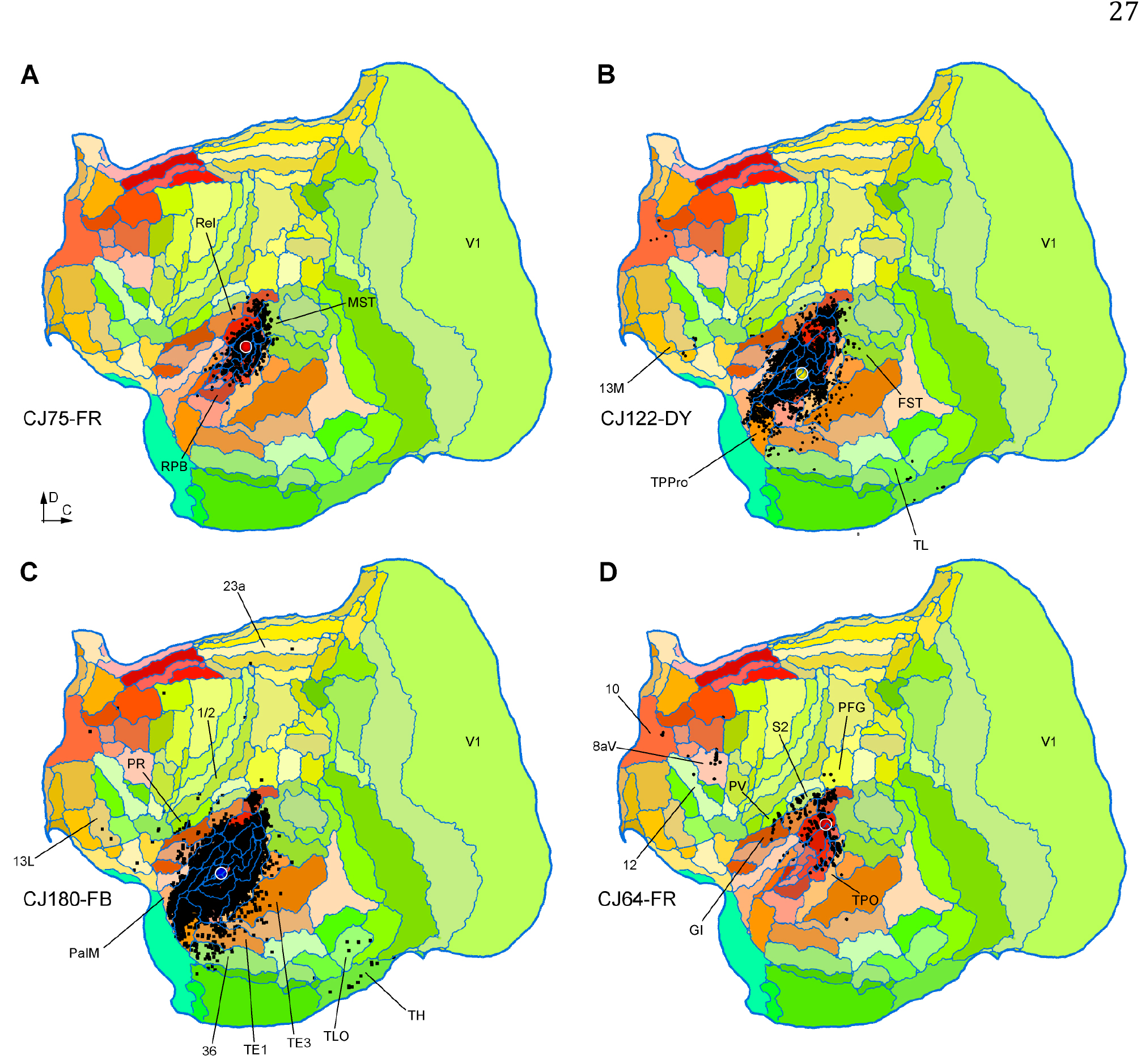
**A-B:** Unfolded reconstructions of the cortex showing the locations of labelled neurons in 4 injections in auditory cortex. The injections illustrated in panels A-C were centred in auditory core areas (with invasion of adjacent areas in B, C), while the one illustrated in D was centred in caudal belt area CM, with likely involvement of A1. None of these injections resulted in retrogradely labelled neurons in V1. Several cortical areas where long-range projection neurons were labelled are indicated for orientation. For abbreviations, see Table 2.

**Figure 9:**
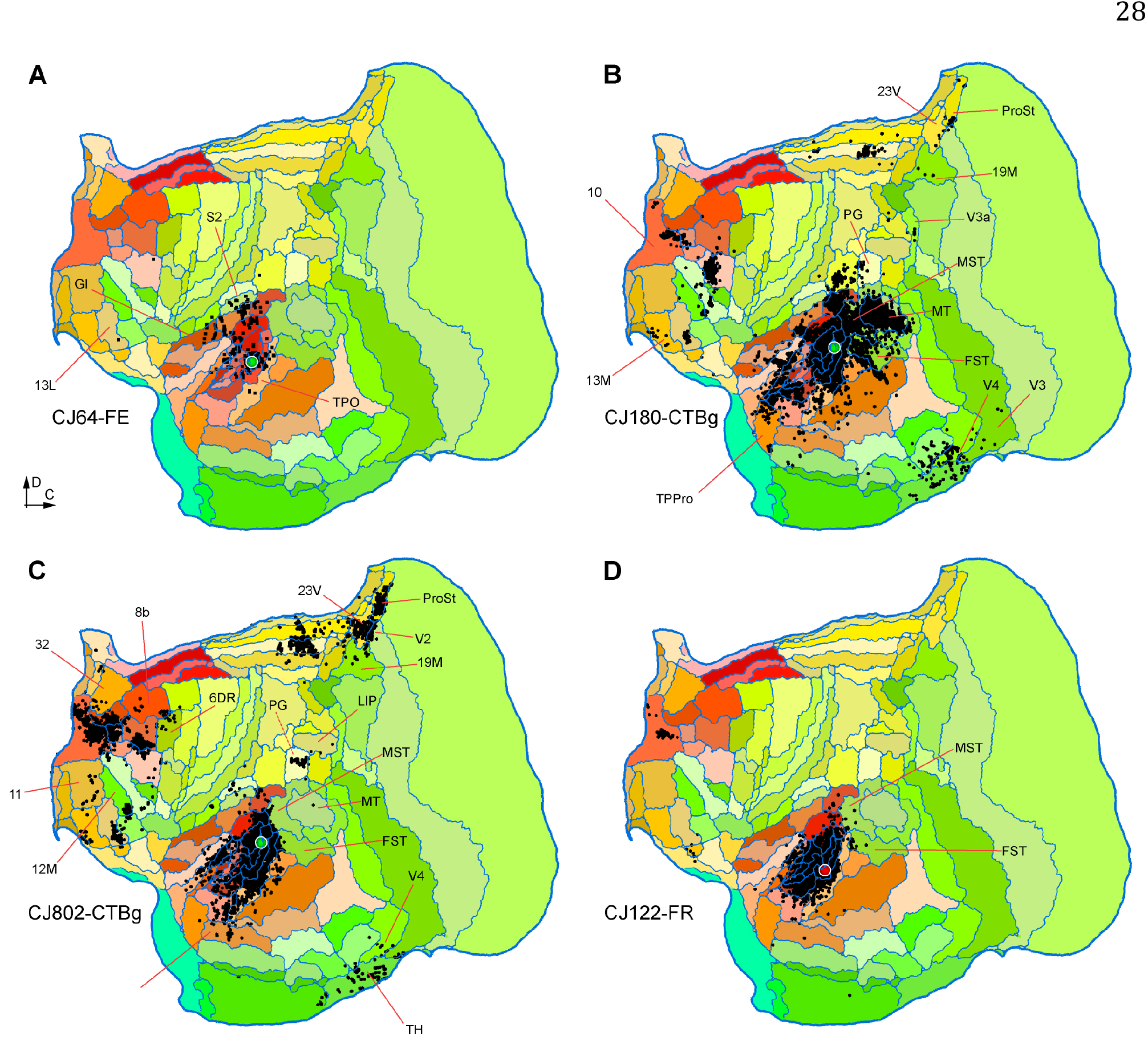
**A-D:** Unfolded reconstructions of the cortex showing the locations of labelled neurons in 4 injections in auditory cortex. A-C are injections involving area ML, and D is an injection centred in the caudal parabelt. Several cortical areas where long-range projection neurons were labelled are indicated for orientation. For abbreviations, see Table 2.

Animal CJ75 had a FR tracer injection that encompassed all layers of the lateral part of A1. As shown in Figure 8A, this injection resulted in a very concentrated pattern of label, mostly involving other auditory areas. The only possible visual projections originated in the most rostral part of the medial superior temporal (MST) and fundus of superior temporal (FST) areas, which accounted for 6.04% of the extrinsic afferents. As discussed below, the rostral sector of MST was the most commonly labelled region among putative visual areas, suggesting that this area, as defined in Paxinos et al. 2012, may be functionally heterogeneous. Putative polysensory projections were observed from the retroinsular (ReI) and temporoparietal transition (TPt) areas.

Two other injections involved the auditory core areas. In case CJ122-DY (Fig. 8B), the injection site was near the border of the rostral (R) and rostrotemporal (RT) areas, with possible involvement of the anterolateral belt area (AL), whereas in case CJ180-FB (Fig. 8C) it was centred in RT, with involvement of the rostrotemporal lateral area (RTL) and AL. Again, putative visual connections originated in the rostral parts of MST and FST, which corresponded to small proportions of the extrinsic label (0.96% and 1.09% in CJ122-DY and CJ180-FB, respectively). However, in these cases additional connections from putative visual association cortex in the inferior temporal cortex (areas TE1-TE3; see Fig. 8C).

Three injections were centred in the caudal belt areas. In case CJ64- FR (Fig. 8D), a small injection near the border of areas CM, CL and A1 failed to label neurons in visual areas, despite clear long-range transport to the frontal lobe, and the presence of cross-modal projections from somatosensory cortex (second somatosensory [S2], parietal ventral [PV] and granular insula [GI] areas). Injections centred in the middle lateral area (ML), with variable involvement of adjacent areas (Table 1), were more heterogeneous in their results (Fig. 9A-C). The most rostral of these injections was a small FE deposit in case CJ64 (Fig. 9A), located right at the boundary between ML and the anterolateral area (AL), involving portions of both areas. In this case, no putative visual afferents were detected. The more caudal injections in ML (cases CJ180- CTBg and CJ802-CTBg; Figs. 9B and 9C) both resulted in label in area prostriata (n=31 and 183 neurons), which is a subdivision of the retrosplenial cortex where visual responses are routinely obtained (Yu et al. 2012), as well as variable amounts of label in areas such as the middle temporal area (MT), MST, FST, the representation of the far periphery of the visual field in V2, and the medial subdivision of area 19 adjacent to peripheral V2 (area 19M; Paxinos et al. 2012), which also predominantly represents peripheral vision (Rosa and Schmid, 1995). Finally, an injection in the caudal parabelt area revealed sparse connections with MST, FST, and inferior temporal areas (9D).

## Discussion

The existence of polysensory responses in macaque cortex has been known for some time, particularly in regions of the superior temporal polysensory (STP) cortex, posterior parietal cortex, and caudal lateral sulcus, which are located at the interface of the classical visual, somatosensory and auditory cortices (Hyvarinen and Shelepin 1979; Bruce et al. 1981; Baylis et al. 1987; Hikosaka et al. 1988; Mazzoni et al. 1996; Schlack et al. 2005). This type of activity was initially thought to be dependent on bottom-up integration of unisensory processing pathways into specialised regions of the cortex, in light of a hierarchical processing model (Jones and Powell 1970; Felleman and Van Essen 1991). However, indications of auditory-induced modulations in the electrophysiological and haemodynamic activity of V1 (Giard and Peronnet, 1999; Macaluso et al., 2000), followed by pioneering studies which revealed that V1 receives direct projections from both the STP cortex and auditory areas (Falchier et al. 2002; Rockland and Ojima 2003) have promoted a revision of multisensory integration models in the cortex. Here we found evidence that cross-modal projections exist in the marmoset. As in the macaque, the projection originates primarily from the infragranular layers of caudal auditory areas. Although they form a small minority of the cortical extrinsic afferent projections to V1, auditory projections appear to be specific. For example, no labelled neurons are located in the rostral auditory cortex, or other cortical areas located at similar or shorter distances from the injection sites. Moreover, in several of our cases labelled neurons were observed in the visual domains of the frontal lobe (areas 8aV and 45; Burman et al. 2006; Reser et al. 2013), demonstrating that the lack of other projections is not the result of short transport times following the injections.

The primary significance of these results is that they confirm that auditory areas send direct, monosynaptic connections to V1 in a New World monkey species. Like macaques, marmosets are simian primates, but, together with other New World monkeys, they have evolved in isolation from Old World monkeys, apes and humans for more than 40 million years (i.e. approximately half of the total evolutionary history of the Order Primates; Perelman et al. 2011; Rosa et al. 2018). This finding suggests that direct auditory influence on early visual processing originated early in primate evolution. Marmosets and macaques differ substantially in terms of body size, brain volume and degree of gyrification, and many behavioural characteristics (Stephan et al. 1981; Rosa et al. 2018). Thus, the preservation of similar auditory projections to V1 seems to suggest a general functional significance across different ecological niches.

There are a few differences in the results obtained in the two species, but these may be due in part to methodological issues. In general, the smaller size of the marmoset brain imposes the use of smaller volumes of tracer, in order to minimise the risk of contamination of adjacent areas or invasion of the white matter. Thus, the numbers of labelled cells in our experiments tend to be much lower than those reported by similar work in the macaque. Considering that the auditory projections to V1 form a relatively low percentage of the total afferent population (<1.0%), the risk of false negatives is increased. In our study, all of the injections that revealed clear projections used the retrograde tracer FB, which tends to form relatively large injection sites, and consequently label larger number of cells (particularly in comparison with dextran-based tracers such as FR and FE). FB was also used in most of the cases reported by Falchier et al. (2002).

In the macaque (Falchier et al. 2002), auditory projections to peripheral V1 appear to be more substantial than those to central vision. This is also apparent in our materials, where the 3 cases with substantial label in auditory cortex (CJ174, CJ178 and M820) all had injections in the parafoveal or peripheral representation. Case M822, where a large FB injection was placed near the representation of the fovea, only revealed a few labelled neurons in auditory cortex, and no label was detected in other cases. However, one of the animals (CJ82) had injections of FR and DY in the peripheral representation (10–20° eccentricity) which resulted in substantial long-range transport, but failed to label auditory areas. In considering these results, it may be relevant that the latter injections were very close to, or straddled the V1/ V2 border, where the vertical meridian of the visual field is represented. In humans, estimates of minimal detectable separation between auditory stimulus sources are highest when the stimuli are arranged vertically along the midline (Perrott and Saberi 1990), and the benefit of concurrent auditory stimulation in enhancing spatial localisation of visual stimuli is not observed near the midline (Perrott et al. 1993). This suggests the interpretation that auditory projections to V1 may be primarily directed to representations of portions of space away from the vertical meridian, where they can be of greatest value in stimulus localisation, rather than to the peripheral representation of this area in general. Other studies have shown that visual stimulus localisation is only enhanced by concurrent auditory stimulation in the peripheral visual field, which is usually defined as distance from the fovea along the horizontal dimension (Perrott et al. 1991; Frassinetti et al. 2002; Gleiss and Kayser 2013), and that the sensation of self-motion is enhanced by concurrent auditory and peripheral visual stimuli simulating vection (Keshavarz et al., 2014). There is also physiological evidence in macaque monkeys that indicates that concurrent auditory stimuli reduces the saccadic reaction times to peripheral visual stimuli (Wang et al. 2008).

Auditory projections to V1 have been well documented in cats (Hall and Lomber 2008), in a study which also found that the projections to the peripheral representation are more substantial. Neuroimaging and electroencephalographic studies have found various forms of modulation of the haemodynamic activity by concurrent audiovisual stimulation, and even evidence of auditory-evoked activity in visual areas (e.g. Giard and Peronnet 1999; Noesselt et al. 2007; Watkins et al. 2007; Brang et al. 2015; Azevedo et al. 2015; Petro et al. 2017), as well as evidence of functional connectivity between unimodal visual and auditory areas (Eckert et al. 2008). However, these modulations may not necessarily depend on monosynaptic connections (Chaplin et al. 2018). In rodents, where the smaller brain and reduced number of areas imposes a different connectional architecture characterised by more direct interconnectivity between systems (Horvát et al. 2016; Gămănuţ et al. 2018), direct connections from auditory cortex to V1 appear to be more common (Larsen et al. 2009; Izraeli et al. 2002), but resemble those present in primates in terms of the infragranular predominance of the neurons forming connections (Charbonneau et al. 2012; Laramée et al. 2013). Unlike in primates, a reciprocal projection from V1 to auditory cortex has been described in prairie voles (Campi et al. 2010).

Using 8 tracer injections, we have explored the possibility of V1 projections to auditory areas. Two of these injections (CJ180- FB; Fig. 8C, and CJ122- DY; Fig. 8B) were relatively large, and involved subdivisions of the rostral core and belt areas. Neither of these resulted in any label in V1, or most other visual areas, despite revealing long-range connections from frontal and parahippocampal areas; thus, we are confident that the rostral auditory areas, which are thought to be involved in identification of sounds (Tian et al. 2001), do not participate in cross-modal connections with areas corresponding to low hierarchical levels of processing in visual cortex. One possible exception is area MST: in both cases, labelled neurons were found in the most rostral part of MST (as defined by myeloarchitecture; Palmer and Rosa 2006a, b). The spatial specificity of this connection could be seen as indication that MST, as currently defined in the marmoset, is heterogeneous. In particular, the marmoset superior temporal polysensory cortex (area TPO) could extends further dorsally than currently estimated, forming a narrow belt of cortex that separates caudal auditory areas CM and CL from MST proper (Fig. 10). Indeed, studies in the macaque have indicated that cytoarchitectural area TPO is heterogeneous (Padberg et al. 2003), and it is possible that a dorsal, more densely myelinated sector, has been confounded with MST in the parcellation proposed by Paxinos et al. (2012). Evaluation of this hypothesis will require tracer injections in different portions of this region. In agreement with observations in the macaque (Hackett et al. 2007), we also found sparse somatosensory inputs to auditory areas, and many of the auditory injections labelled neurons in parahippocampal cortex, where sensory convergence has also been documented (Blatt et al. 2003).

**Figure 10:**
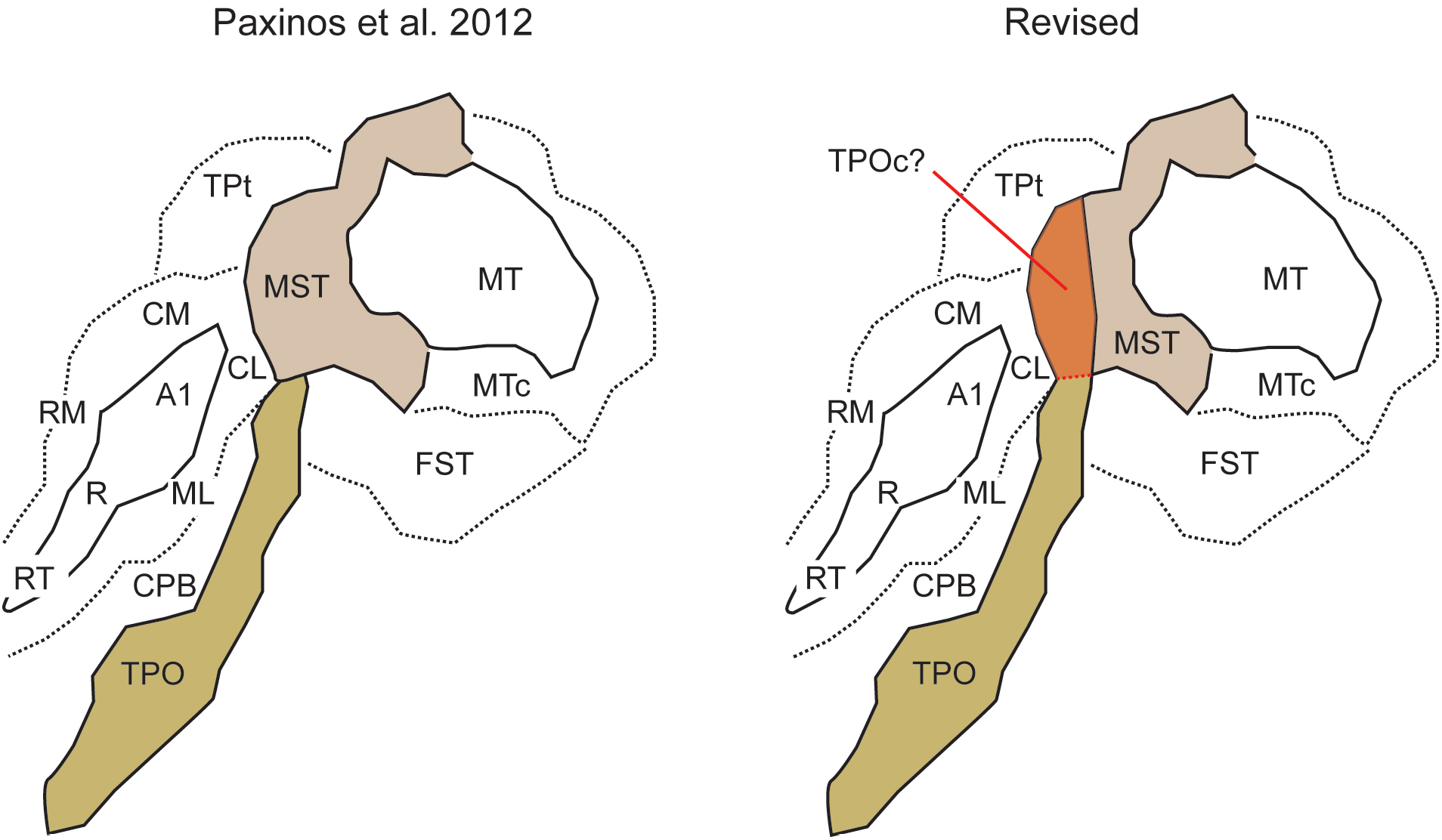
A proposal for parcellation of marmoset area MST in two sectors, based on the present data and comparison with results obtained in the macaque monkey (Falchier et al. 2002). **Left:** original parcellation of the cortex near the tip of the superior temporal sulcus, according to the architectural study of Paxinos et al. (2012). The diagram follows the orientation of Figure 2. **Right:** The sector of MST where most labelled neurons are observed following auditory cortex injections could be a homologue of the caudal subdivision of area TPO (TPOc;Padberg et al. 2003), bringing in line the results obtained in macaque and marmoset monkeys.

The only clear cross-modal visual projections observed to auditory cortex in our data targeted auditory belt area ML. Both of the injections in this area (Figs. 9B, C) revealed a substantial projection from area prostriata, a subdivision of visual cortex characterised by large, peripheral receptive fields, and very short latencies to visual simulation (Yu et al. 2012). In the macaque, Falchier et al (2010) also reported projections from prostriata to other subdivisions of caudal belt auditory cortex, suggesting that this projection targets multiple areas involved in auditory localisation. One of the injections in area ML labelled, in addition, substantial numbers of cells in area MT, and both injections resulted in scattered label in the peripheral representations of areas V2, V3 and V4. As a whole, the peripheral field specificity of these projections, together with the likely involvement of caudal auditory areas in spatial aspects of hearing (Tian et al. 2001) suggest that information from the far periphery of the visual field may modulate auditory processing, probably enhancing detection and localisation of stimuli.

In summary, our data add to the growing evidence pointing to the fact that, in addition to the bottom-up integration of sensory signals across hierarchical pathways converging into polysensory association areas, the primate brain contains sparse, but direct and specific pathways that allow information flow between areas traditionally conceived as unisensory. The auditory connections to V1 and MT (present results; Palmer and Rosa 2006a) are likely to play roles in enhancing stimulus detection and localisation, particularly away from the vertical meridian. Visual projections to auditory areas, primarily from prostriata and other peripheral representations, may provide a complementary role in detection, especially in situations when the signal to noise ratio of the auditory stimulus is low.

## Ethical approval

All procedures performed in studies using animals were in accordance with the ethical standards of the Institutions (Monash University and the RIKEN) at which the studies were conducted. All applicable international, national and institutional guidelines for the care and use of animals were followed.

## Funding

- Australian Research Council. Grant numbers: DP140101968, CE140100007
- International Neuroinformatics Coordinating Facility Seed Funding Grant
- Japan Agency for Medical Research and Development. Grant number: JP17dm0207001 (Brain/MINDS)

## Conflict of interest

The authors declare that they have no conflict of interest.

